# Porous upconversion nanostructures as bimodal biomedical imaging contrast agents

**DOI:** 10.1101/837864

**Authors:** Ziqing Du, Abhishek Gupta, Christian Clarke, Matt Cappadana, David Clases, Deming Liu, Zhuoqing Yang, Philip Doble, Bill Price, Xiaoxue Xu

## Abstract

Lanthanide ions doped upconversion nanoparticles (UCNPs) hold great promise as the imaging contrast agent for multimodal medical imaging techniques for diagnosis, including fluorescent bioimaging, magnetic resonance imaging, and computed tomography. However, the maximized signal values of fluorescence and MRI cannot be achieved simultaneously from the same upconversion nanoparticles structures because high specific surface areas can benefit the signal gaining of MRI while big size can induce brighter fluorescent imaging. In this work, we designed and fabricated novel core-porous shell structures for UCNPs with much-enhanced signal values for both fluorescent imaging and MRI. The core-porous shell UCNPs were synthesized via a post-treatment process after an inert shell was coated onto the core UCNPs. The formation mechanism was carefully investigated. The fluorescent and magnetic resonance properties have been detailed characterized and compared from core, core-shell and core-porous UCNPs. Large and bright UCNPs in fluorescence and MRI have been achieved and great potential as the dual-modal contrast agent.

---

A wide variety of non-invasive bioimaging techniques are commonly employed for clinical diagnosis and scientific research, including ultrasound, computed X-ray tomography (CT), magnetic resonance imaging (MRI), positron emission tomography (PET), and bioluminescence imaging (BLI) [1–5]. Correspondingly, various exogenous materials as contrast agents are adopted for specific techniques to enhance signal-to-noise ratio [6]. For instance, nano-bubbles can enhance the ultrasound scanning resolution [4, 7–9], Iodine-based and barium-sulfate based material with strong X-ray absorption are used for CT and PET [6, 10], and magnetic materials such as iron oxide or gadolinium compounds are best for MRI to gain paramagnetic signals [11, 12]. Fluorescent dyes, such as firefly luciferase (FLuc), is benefited by very high sensitivity in bioluminescence imaging and a broad dynamic range relative to other imaging techniques. [13, 14].

Multimodal contrast agents that can be adapted to more than one bioimaging techniques [15] are highly advantageous for obtaining enhanced accuracy and sensitivity imagery [11, 16] and improved patient care [5]. Bioimaging modalities can complement with each other for a more thorough examinations. For example, optical fluorescent microscopy can achieve single molecular resolution in *in vitro* imaging[13, 14, 17], while CT is benefited on the hard tissue detection and 3D imaging with high reliability and rich details [18, 19], and MRI presents higher resolution images on muscle and organs [20, 21]. Furthermore, fewer possible side effects can be resulted from the application of multimodal contrast agents, due to its relative smaller necessary dose and ability of serving multi technologies [11, 22].

Lanthanide ions (Ln) doped upconversion nanoparticles (UCNPs) have been demonstrated as a robust multimodal contrast agent owing to a variety of magnetic and optical properties arising from 4f electrons in Ln^3+^[23, 24]. UCNPs show tremendous potential in deep-tissue optical bio-imaging, MRI, CT and PET scan for the guidance of high precision surgery [23, 25]. Lin’s group [26] had firstly reported the dual-modality imaging, MRI, and luminescent imaging of living whole-body animal using Tm^3+^/Er^3+^/Yb^3+^ co-doped NaGdF_4_ UCNPs. Van Veggel et al. [27] presented 200-3000 times higher ion relaxivity per contrast agent from 2.5 nm ultrasmall NaGdF_4_ UCNPs than the clinical agent. Their work also revealed that the surface-to-volume ratio of the UCNPs was the major contributor to the relaxivity value, so the smaller the UCNPs possessed higher T1 relaxivity. However, own to their higher surface quenching, the luminescent emission intensity would not be satisfied with ultrasmall UCNPs. Thus, a core-shell structure was used for dual-modality imaging. Capobianco et al. [28] further confirmed that the ultrasmall NaGdF_4_ in 5 nm UCNPs presented high relaxivities and strong vascular signal enhancement compared to 25 nm one as dual MR/optical imaging probe. Yang et al. [29] demonstrated the three imaging modalities of the core-shell NaGdF_4_:Yb^3+^, Er^3+^-NaGdF_4_:Nd^3+^UCNPs in MR/optical/CT techniques. The obtained T1 relaxivity 5.7 mM^-1^ s^-1^ at 3T magnetic field from 26 nm core-shell UNCPs was comparable to the Gd-DTPA in clinic. Another core-shell-shell UCNP nanostructure in 35 nm were demonstrated with multiple imaging capabilities in UCL/CT/MRI [30]. Feng et al. [31] adopted a new host crystal of Ba2GdF7 doped with Yb^3+^ and Er^3+^ for bright UCL and reasonably high CT/MR. Heterogeneous core-shell NaGdF_4_@CaCO_3_ nanoparticles have been fabricated for the MRI and ultrasonic dual-modal imaging [32]. Despite the excellent performance of core-shell UCNPs in luminescence imaging, there is a dilemma between the optical emission intensity and the MRI/CT signals for UCNPs, which can be attributed to the size of the nanoparticles. The host-guest system of UCNPs requires bigger size for high emission intensity, while UNCPs doped with Gd ions needs higher specific surface areas to increase MRI signals [33, 34].

In this work, we proposed a novel UCNPs in a core-porous shell structure with well-maintained luminescent emission intensity and much enhanced MRI signals. The formation mechanism is illustrated in Figure 1. Active and bright UCNP core would be fabricated with relatively bigger size, protective shell containing Gd^3+^ will be coated with tuneable thickness, later the protective shell will be post-treated at elevated temperature in acidic environment to porous structure to increase the exposure surface areas of Gd^3+^. In the core-porous shell nanoparticles, active NaYF_4_ nanoparticles doped with Yb and Er ions responsible for high emissions for luminescent imaging and the exposed Gd^3+^ in the porous shell increase the contact areas to protons for higher MRI signals.

**Figure 1.**
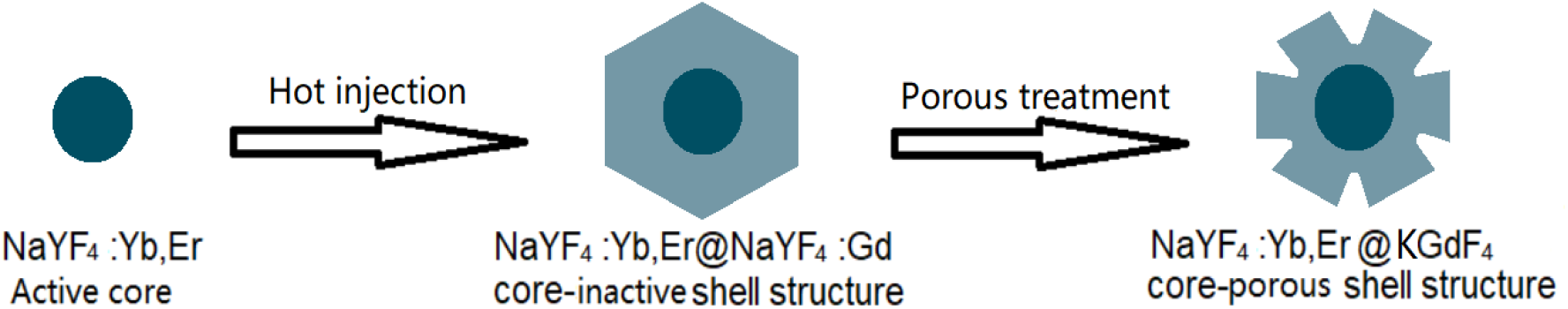
Design scheme for core, core-shell and core-porous shell UCNPs.

We fabricated the core, core-shell and core-porous shell UCNPs and conducted characterizations on morphology. From Figure 2 we can observe the evolution from core, core-shell and core-porous UNCPs with the core size of 22.5 nm (Figure 2 (a-d)) and 36.5 nm (Figure 2(e-h)). Figure 2(a) presents the highly uniform core UNCPs of NaYF_4_: 20%Yb, 1%Er in the size of 22.5 nm. The core-shell UNCPs of NaYF_4_: 20%Yb, 1%Er @ NaYF_4_: 30%Gd was obtained with uniform shell coating in the thickness of 6 nm. The elemental composition of the core-shell UNCPs are shown in Figure S1(a) and it clearly shows that Gd^3+^ only was contained in the shell. After the post treatment in a thermal-acidic environment to the core-shell UCNPs, the porous shell was formed as shown in Figure 2(c). The flower-shaped surface displayed the connected 3D structure, suggesting the selective etching occurred on specific spots. The high-resolution image was shown in Figure S1(b). The crystallinity was well maintained, and crystal orientation was remained the same with the core-shell UCNPs (more details are required). The elemental composition for the core-porous shell UNCPs is shown in Figure 1(d). It can be seen that the Y^3+^ distributes more in the core area and the Gd^3+^ ions’ distribution is not continuous and exhibits voids demonstrating the porous shell.

**Figure 2.**
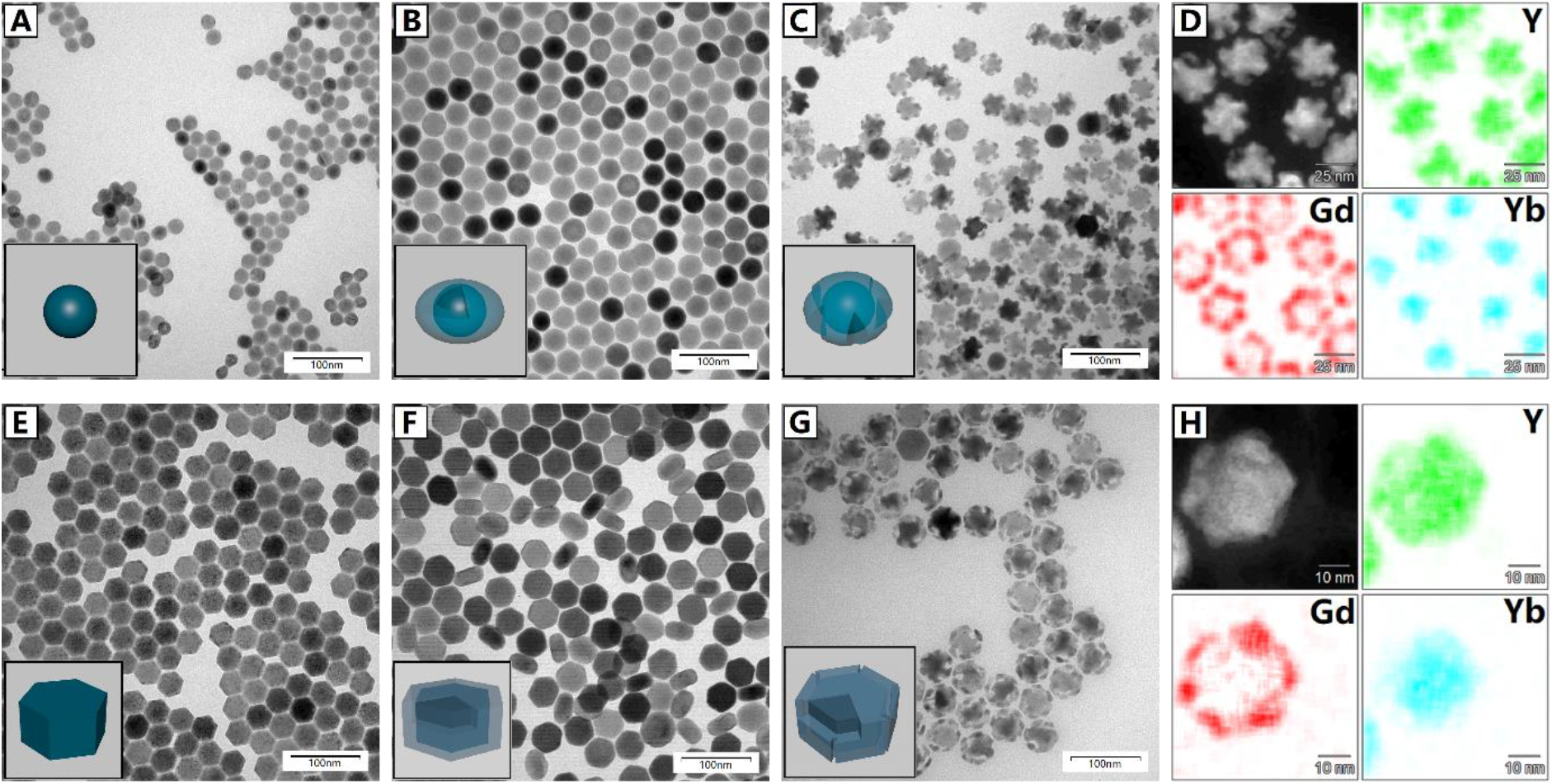
TEM images, and composition analysis of NaYF_4_: 20%Yb, 1%Er bare nanoparticles with d = 22.5nm (A) and 36.5nm (E), NaYF_4_:20%Yb,1%Er@NaYF_4_:30%Gd core-shell nanoparticles with shell thickness 6nm (B, F), and NaYF_4_: 20%Yb, 1%Er @ NaYF_4_: 30%Gd @ K_2_Y_x_Gd_1-x_F_5_ core-porous shell nanoparticles (C, G) under 18000 time magnification (scale bar equals 100nm). The EDS result shows the distribution of Y, Gd and Yb ions on core-porous shell nanoparticles (D, H). The size distribution result of all samples contained in supporting information.

Similarly, the 6 nm thick dense shell and porous shell can be can be fabricated onto the bigger core UCNPs (d = 36.5nm) showing in Figure 2(e-g). The porous shell is a bit different from that in the Figure 1(c) because the etching extent was controlled for a shorter time. The frame of the shell was maintained quite well, the etching occurred more deeply inside the shell. The elemental mapping images also shown relatively completed shell by Gd^3+^ and Y^3+^.

The full energy-dispersive X-ray spectroscopy (EDX) spectra for the core-shell and core-porous shell UCNPs with two sized cores were shown in Figure S1(d). The potassium ions K^+^ was detected and presented within the porous shell, indicating the K_2_Y_x_Gd_1-x_F_5_ might be formed while the post treatment in a slight amount in a slight amount. The crystal phase of newly formed structure was not detectable using XRD and the result was shown in Figure S2. The hexagonal phase of all NaYF_4_ samples were confirmed using XRD. The phases of the UCNPs were still hexagonal without observable changes after the shell coating and the post-treatment.

With the confirmation in phase and morphology for the UCNPs, we conducted the photoluminescent measurements for the core, core-shell and core-porous shell UCNPs in dispersions. The spectra were shown in Figure 3(A&B) for the 22.5 nm and 36.5 nm core, and corresponding core-shell and core-porous shell UCNPs respectively. The batch of the core, core-shell and core-porous shell UCNPs with bigger diameter of cores presented much higher emission intensity than the batch UCNPs with small cores. For both series, the core UCNPs display lowest emission intensities at 524 nm, 545 nm, and 675 nm, while the emission intensities from the dense shell coated UNCPs were much enhanced. The enhancement factor was 2.5 and 10 For the UCNPs with 22.5nm and 36.5nm core with 6 nm dense shell respectively. The emission intensity of the core-porous shell UCNPs decreased for both batches compared to UCNPs with dense shell, but the enhancements were still quite significant compared to the bare core and the enhancement factors were roughly 1.5 and 9. It is understandable because the inert shell plays an important role for emission improvement due to the minimization of the surface quenching, and it is known that the thicker the shell is up to 10 nm, the better it protects the core [35]. When the inert shell was etched, the protection effect is deteriorated slightly while the UCNPs were still bright enough as the fluorescent imaging probe.

**Figure 3.**
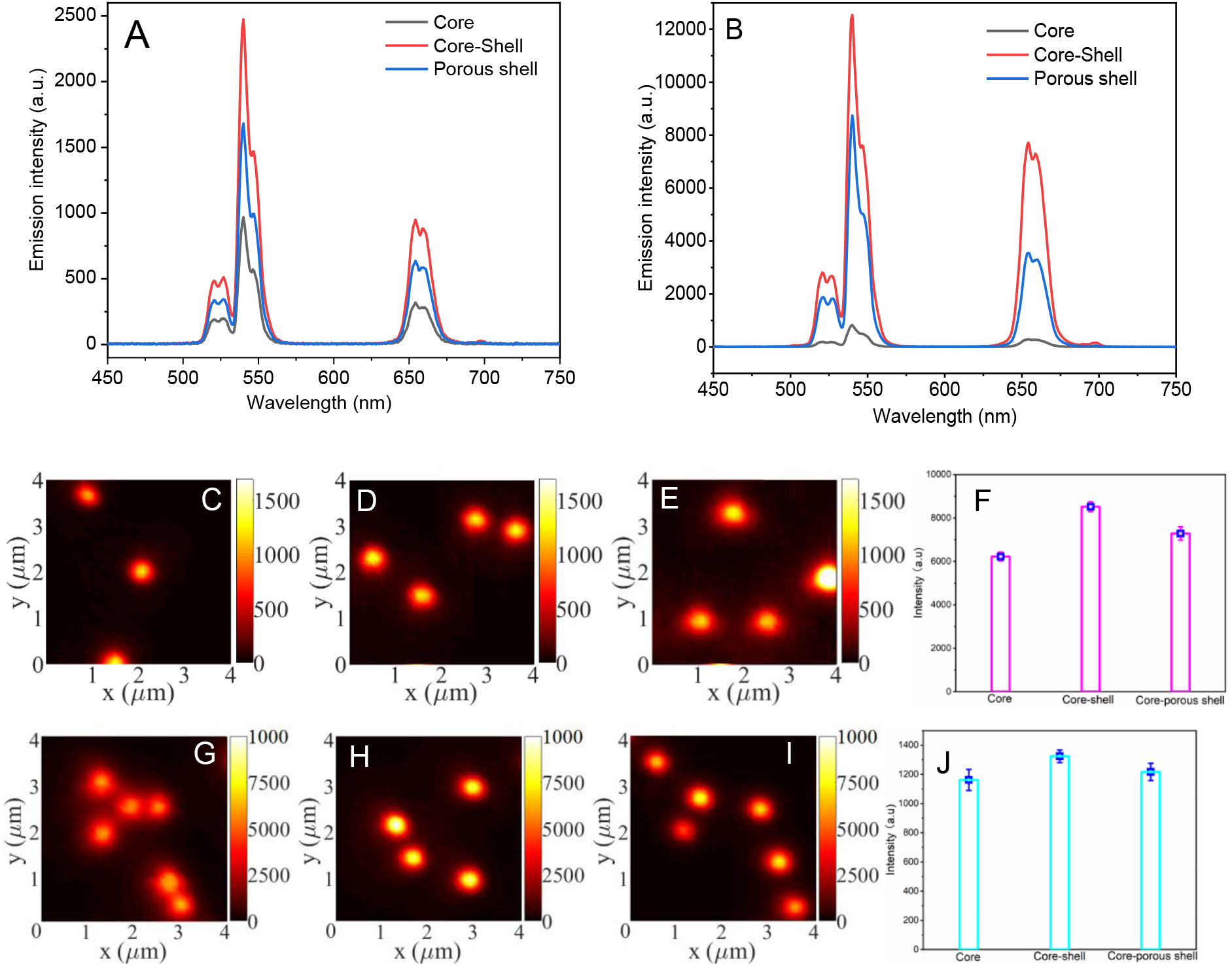
Photoluminescence spectroscopy of core, core-shell and core-porous shell nanoparticles with core diameter equals 22.5nm (A) and 36.5nm (B). All analysis was processed under power of 2.723 Watts. Single Particle fluorescence images of (C, G) bare core nanoparticles with different diameters; (D, H) core-shell nanoparticles; (E, I) core-porous shell nanoparticles. The quantitative comparisons were shown in (F, J).

We further carried out the single nanoparticle characterization for the two batches UCNPs using a homebuilt scanning confocal microscopy. The single particle fluorescence imaging results in Figure 3 (C-J) agree well with the PL measurements for the dispersion UCNPs. The core, core-shell and core-porous shell UNCPs batch with big core exhibited much higher emission intensity than the batch with small core, which are consistent with the intensity of the UCNPs dispersions. The emission intensity of the UCNPs was enhanced greatly and then dropped a bit with the dense shell growing and porous treatment. But it can be seen that the enhancement factors in the single nanoparticles of both batches were not as significant as that for the bulk dispersion samples. The enhancement factors are 1.17 and 1.04, 1.38 and 1.08 from the core-shell and core-porous shell UCNPs with small and big core UCNPs, own to the surface quenching effect from the environment to the core in single UCNPs is much less than the UCNPs in dispersions.

To verify the second imaging modality of the core-shell and core-porous UCNPs, we carried out the NMR measurements for both the batches. It can tell from the comparison results in Figure 4 that the porous-shell UCNPs displace much higher r_1_ and r_2_ values than the core-shell UCNPs at both 1 Tand 11.7 T magnetic fields. The bigger UCNPs batch exhibits higher r_1_ and r_2_ values than the small UCNPs batch. The comparison from our work to other reports is summarized in table S1.

**Figure 4.**
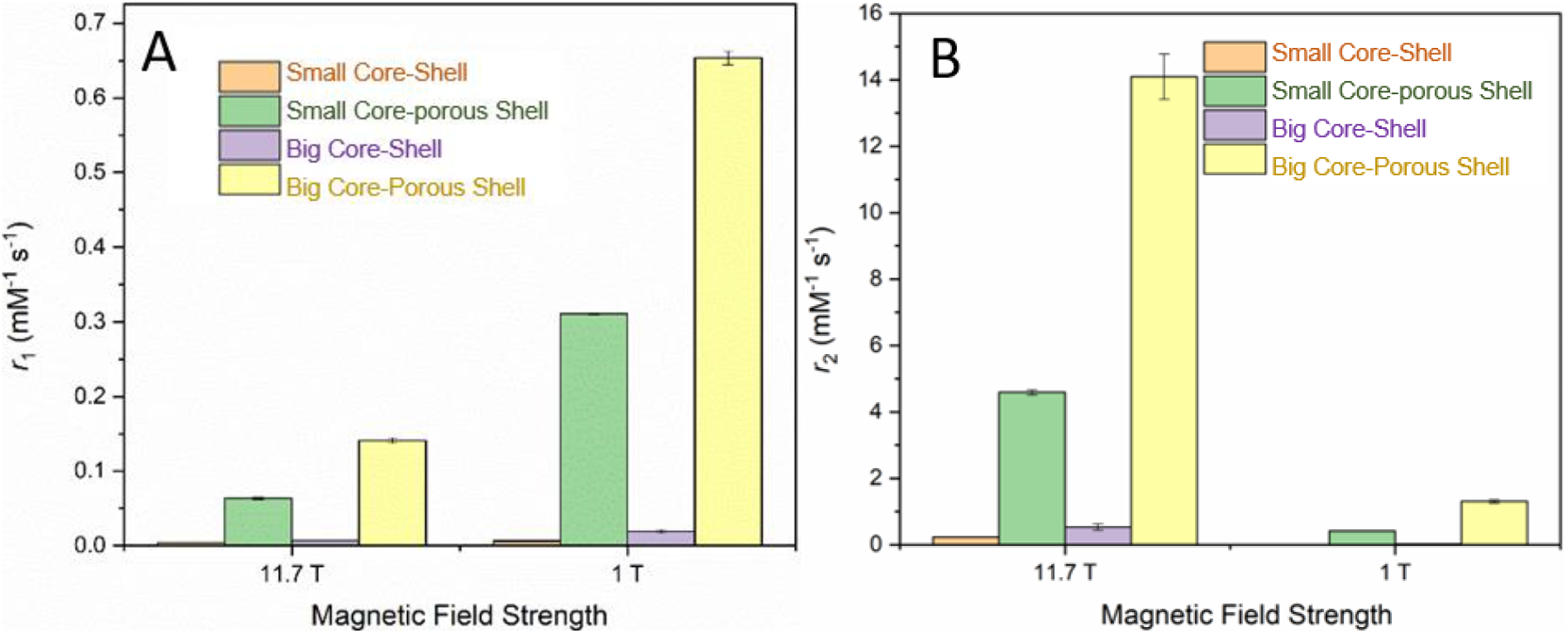
The magnetic relaxivity of r_1_ (A) and r_2_ (B) for the core, core-shell and core-porous shell UCNPs at 1T and 9.4T.

We have designed and fabricated the novel core-porous shell UCNPs with different sized core nanocrystals. An inert and dense shell was coated onto the core UCNPs and the fluorescence intensity was much improved. The controlled etching treatment to the dense shell containing Gd^3+^ and Y^3+^ ions of the core-shell UCNPs induced porous structure and much high exposure of the Gd^3+^, thus the MRI signal values was enhanced greatly. Brighter UCNPs were achieved in both fluorescent and magnetic resonance imaging simultaneously. The core-porous UCNPs can be used as contrast agent in dual modality of biomedical fluorescent imaging and magnetic resonance imaging for diseases diagnosis.

## Acknowledgements

We thank Dr Chao Shen for the great help at TEM characterization in the Microscopy Unit at Macquarie University as well as Microscopy Analysis Unit at the University of Technology Sydney. We would like to thank the Biomedical Magnetic Resonance Imaging Facility at the University of Western Sydney node of the National Imaging Facility (NIF). This project was primarily supported by the Chancellor Postdoctoral Research Fellowship of University of Technology Sydney (Xiaoxue Xu).

